# Drosophila Pyruvate Kinase Links Metabolic State with Circadian Output via TARANIS and PDF

**DOI:** 10.1101/2025.08.18.670854

**Authors:** Sang Hyuk Lee, Oghenerukevwe Akpoghiran, Eunjoo Cho, So Who Kang, Min-Ji Kang, Kyunghee Koh, Eun Young Kim

## Abstract

The circadian clock generates ∼24-hour rhythms that anticipate daily environmental changes. Circadian clock and glucose metabolism are tightly interconnected, and both are disrupted in aging and disease. To examine how glucose hypometabolism impacts circadian rhythm, we downregulated glycolytic enzymes - *Hexokinase*-*C* (*Hex*-*C*), *Phosphofructokinase* (*Pfk*), and *Pyruvate kinase* (*Pyk*) - in *Drosophila* clock cells. Only *Hex-C* and *Pyk* knock-down (KD) altered period, lengthening and shortening rhythms, respectively. Notably, *Pyk* KD induced period shortening persisted in adult-specific KD (AKD), indicating a role independent of developmental effects. *Pyk* AKD reduced both PERIOD and Pigment-dispersing factor (PDF) protein levels, with PDF loss driving the short-period phenotype. Mechanistically, the transcriptional co-regulator TARANIS (TARA) was required: *Pyk* AKD lowered *tara* expression, while *tara* overexpression rescued PDF and circadian period. Our findings identify a novel PYK-TARA-PDF regulatory axis linking glycolytic activity to circadian neuropeptide output, providing mechanistic insight into how metabolic dysfunction contributes to circadian disruption in aging and neurodegenerative diseases.

## Introduction

The circadian clock system enables living organisms to anticipate environmental fluctuations driven by the Earth’s rotation, leading to approximately 24h rhythms in behavior and physiology. In animals, this system comprises a cell-autonomous circadian clock, with a master clock located in the brain and peripheral clocks distributed throughout the body (Schibler *et al*, 2015). Primarily entrained by light, the master clock synchronizes the peripheral clocks via nervous and endocrine systems, ensuring the timely coordination of physiological processes and behavior (Mohawk *et al*, 2012). Dysregulation of circadian rhythms is commonly observed in aging and disease conditions, contributing to functional decline and disease progression (Hood & Amir, 2017b; Leng *et al*, 2019).

The mechanism of cell-autonomous circadian clock is highly conserved across animal phyla, relying primarily on transcription/translation feedback regulation of core clock proteins (Patke *et al*, 2020). In *Drosophila*, this regulation involves the transcription factors CLOCK (CLK) and CYCLE (CYC), which activate the transcription of *period* (*per*) and *timeless* (*tim*) genes. The resulting PER and TIM proteins then inhibit the activity of CLK/CYC, initiating the next cycle after being degraded. Additionally, CLK and CYC stimulate the expression of *Par Domain Protein 1ε* (*pdp1ε*) and *vrille* (*vri*) which respectively activate and repress the expression of *Clk*, thereby contributing to the robust organization of feedback loops. *Clockwork orange* (*CWO*), transcribed by CLK/CYC, amplifies the amplitude of CLK/CYC-dependent transcriptional activation and repression cycle (Kadener *et al*, 2007; Lim *et al*, 2007; Matsumoto *et al*, 2007). Homologs and functional counterparts operate in the mammalian circadian system (Patke *et al*., 2020).

In the *Drosophila* brain, approximately 150 clock neurons are organized into several groups based on their anatomical regions, including small ventral lateral neurons (sLN_v_s), large lateral ventral neurons (lLN_v_s), lateral dorsal neurons (LN_d_s), dorsal neurons 1 (DN_1_s), dorsal neurons 2 (DN_2_s), and dorsal neurons 3 (DN_3_s), collectively forming the circadian clock neuronal network (CCNN) (King & Sehgal, 2020)). This network is analogous to the suprachiasmatic nucleus (SCN), the master clock in mammalian brain, and controls various aspects of circadian behavior and physiology. Among these neurons, sLN_v_s play a crucial role in synchronizing the CCNN under constant dark conditions by releasing Pigment-dispersing factor (PDF), a neuropeptide homologous to vasoactive intestinal polypeptide (VIP) in the mammalian SCN (Herzog, 2007).

Each day in most organisms is divided into active/feeding and resting/fasting periods. To maintain homeostasis, diverse metabolic processes including glucose metabolism, are under circadian clock regulation. For instance, anabolic/catabolic reactions such as gluconeogenesis and glycolysis are temporally regulated (Marcheva *et al*, 2013). Metabolic outputs, in turn, provide feedback to the circadian clock system (Asher *et al*, 2008; Cho *et al*, 2019; Lamia *et al*, 2009; Lee *et al*, 2021; Linford *et al*, 2012; Nakahata *et al*, 2008; Rutter *et al*, 2001). Thus, the circadian clock and glucose metabolism tightly regulate each other for organismal health, with dysregulation of these systems frequently co-occurring in disease conditions. Brain hypometabolism, characterized by reduced glucose consumption, is commonly observed in various neurodegenerative diseases and with aging (Camandola & Mattson, 2017; Gonzalez-Redondo *et al*, 2014; Mosconi *et al*, 2008; Zilberter & Zilberter, 2017). Intriguingly, dysregulated circadian rhythms, such as disruptions in the sleep/wake cycle and temperature rhythms, are also often associated with neurodegenerative diseases and aging (Hood & Amir, 2017a; Lananna & Musiek, 2020; Leng *et al*., 2019) and exacerbate their progression (Holth *et al*, 2019; Kang *et al*, 2009; Kress *et al*, 2018; Liu *et al*, 2024; Majcin Dorcikova *et al*, 2023; Tranah *et al*, 2011). Despite numerous observations of co-occurrence of hypometabolic states and dysregulated circadian rhythms, there has been limited direct investigation into the impact of glucose hypometabolism on circadian rhythm behavior.

To investigate how glucose hypometabolism influences circadian timekeeping, we selectively downregulated three rate-limiting glycolytic enzymes - *HexC*, *Pfk,* and *Pyk* - specifically within *Drosophila* clock cells. Among them, *Pyk* KD consistently shortened the circadian period, particularly under adult-specific KD (AKD) conditions that minimize developmental confounds. This period shortening was accompanied by reduced levels of the neuropeptide PDF, indicating that PYK modulates circadian timing through neuropeptidergic output regulation. We also found that the transcriptional coregulator TARANIS (TARA), previously shown to enhance VRI activity (Akpoghiran *et al*, 2024), serves as a key intermediary in PYK’s regulation of *Pdf* expression and period control. These findings reveal a novel PYK-TARA-PDF pathway connecting metabolic perturbation to circadian disfunction, offering mechanistic insights relevant to understanding circadian abnormalities in aging and neurodegeneration.

## Results

### Knockdown of glycolysis rate-limiting enzymes alters circadian rhythm behavior in *Drosophila*

To investigate how glucose metabolism disruption affects circadian rhythm in *Drosophila*, we downregulated key glycolytic enzymes specifically in clock cells (Fig. 1A). Using the *timeless* (UAS)-gal4 (TUG) driver, we expressed RNAi transgenes targeting *Hex-A, Hex-C*, *Pfk* and *Pyk*, which catalyze the irreversible steps of glycolysis. *Drosophila melanogaster* possesses three hexokinase genes (*Hex*-*A*, *Hex*-*C* and *Hex*-*T*), but since *Hex*-*T* is testis-specific, we focused on *Hex*-*A* and *Hex*-*C* (Duvernell & Eanes, 2000; Moser *et al*, 1980). To minimize off-target effects, we employed multiple UAS-RNAi lines for each gene. Two independent *Hex*-*C* RNAi lines (35337, 35338) and two *Pfk* RNAi lines (34336, 105666) exhibited prolonged periods with reduced rhythmicity, while two *Pyk* RNAi lines (35218, 49533) showed shortened periods with reduced rhythmicity compared to controls (Fig. 1B and 1C). qRT-PCR validation confirmed mRNA downregulation in *Hex-C* and *Pyk* KD flies consistent with their period alterations, but not in *Pfk* KD flies, suggesting possible off-target effects (Fig. 1D).

**Figure 1.**
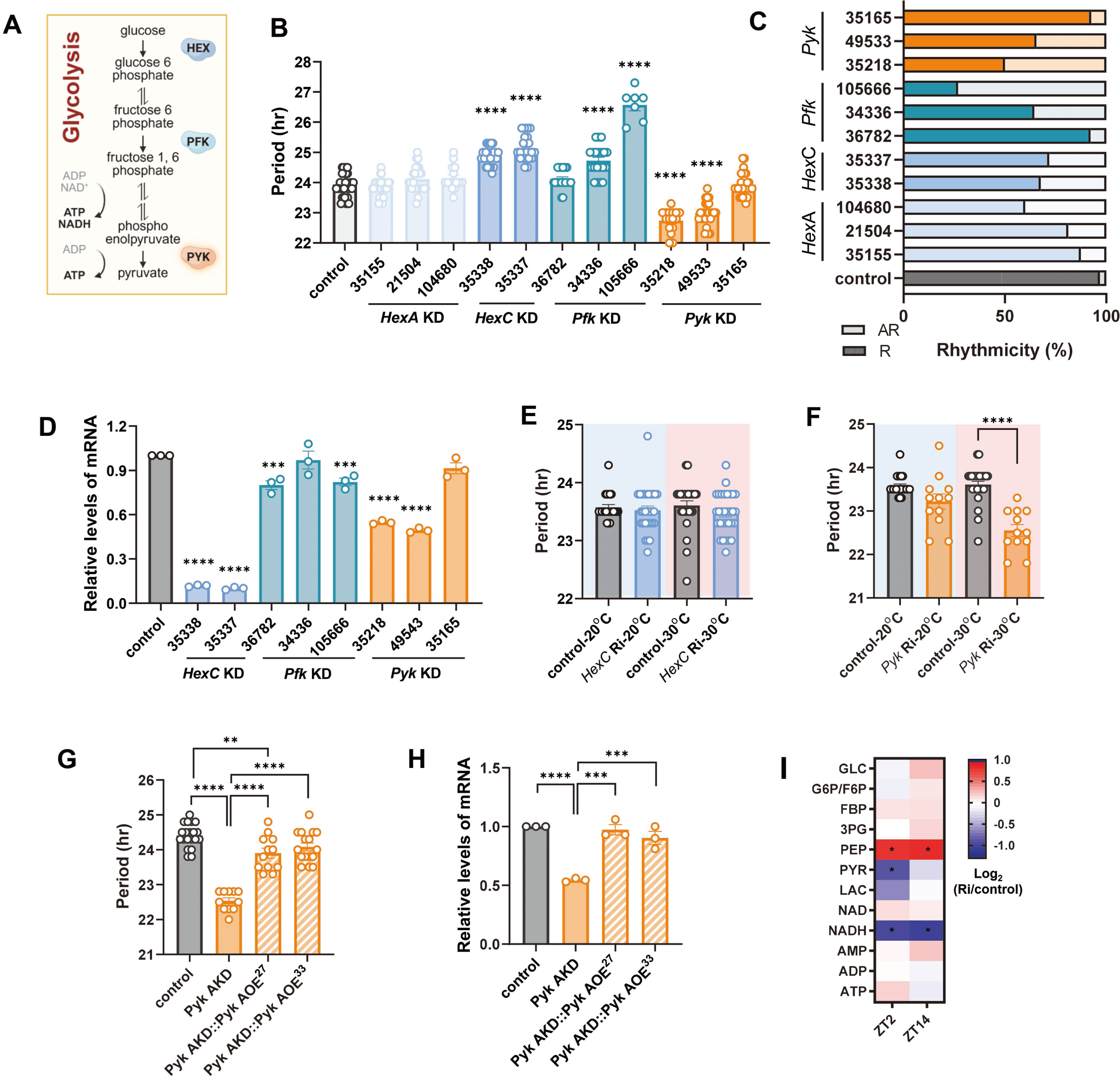
Knockdown of glycolytic enzymes altered circadian rhythm in *Drosophila*. (A) Simplified glycolysis process highlighting the steps catalyzed by *Hex*, *Pfk* and *Pyk.* (B and C) *dcr2*;TUG flies were crossed with *w*^1118^ (control) or indicated RNAi flies. The circadian rhythmic behavior of F1 offspring flies were analyzed at 25°C. (B) Free-running periods of indicated genotype of flies were shown. Bars indicate the mean ± SEM (n = 7–31). Asterisks indicate differences vs. control (one-way ANOVA with Dunnett’s post-test): \*\*\*\**p* < 0.0001. (C) The percentage of rhythmic flies of each genotype is shown. (D) Fly heads were collected at ZT2. mRNA levels of *HexC*, *Pfk*, and *Pyk* were measured by real-time qRT-PCR in control flies and the indicated RNAi lines. Expression levels in KD flies were normalized to those of control flies. Bars indicate mean ± SEM from three independent experiments. Asterisks indicate differences vs. control (*t*-test): \**p* < 0.05, \*\*\**p* < 0.001, \*\*\*\**p* < 0.0001. (E, F) *dcr2*;tub-gal80^ts^;TUG flies were crossed with *w*^1118^ (control) or indicated RNAi flies. F1 offspring flies were reared at 20°C, and the circadian rhythmic behavior was analyzed either at 20°C (blue background) or 30°C (red background). Free-running periods of indicated genotype of flies are shown. Bars indicate the mean ± SEM (n = 12–31), and statistical differences were determined using one-way ANOVA with Tukey’s test: \*\*\*\**p* < 0.0001. (G) Recombinant fly lines bearing *Pyk* Ri and UAS-*Pyk*-HA were crossed with *dcr2*;tub-gal80^ts^;TUG. F1 offspring were reared at 20°C and then shifted to 30°C to activate adult-specific knockdown (AKD) or over-expression (AOE). Free-running periods of indicated genotype of flies were shown. Bars indicate the mean ± SEM (n = 12-20). Statistical differences were determined using one-way ANOVA with Tukey’s test: \*\**p* < 0.01, \*\*\*\**p* < 0.0001. (H) Fly heads were collected at ZT2. mRNA levels were measured by real-time qRT-PCR in control and indicated genotype. Expression levels in genetically manipulated flies were normalized to those of control flies. Bars indicate mean ± SEM from three independent experiments and statistical differences were determined using one-way ANOVA with Tukey’s test: \*\*\**p* < 0.001, \*\*\*\**p* < 0.0001. (I) Control and *Pyk* AKD flies were maintained on a 12L:12D cycle at 30°C. Fly heads were collected at ZT2 and ZT14 on day 7 and processed for metabolomics analysis. Heat map represents log_2_ fold changes of glycolytic metabolites in *Pyk* AKD flies relative to controls. Fold changes and statistical significance were calculated from mean ± SEM values across five independent experiments. Statistical differences were determined using Mann–Whitney U test: \**p* < 0.05.

Given the pivotal role of *HexC* and *Pyk* in glycolysis, continuous KD throughout development might interfere with neuronal network formation, indirectly affecting circadian behavior. To test whether *HexC* and *Pyk* is required in the adult circadian clock, we employed a temperature-sensitive tubulin-Gal80^ts^ (tub-g80^ts^) repressor with the TUG driver to conditionally downregulated these enzymes during the behavioral analysis of adult flies (McGuire *et al*, 2004). Flies reared at 20°C, permissive temperature suppressing RNAi showed no period changes for either knockdown (Fig. 1E). At 30°C, where Gal80^ts^ was inactive and RNAi was induced, *Hex*C AKD flies no significant period change, while *Pyk* AKD flies exhibited shortened periods, consistent with continuous KD results (Fig. 1B and 1F). These results demonstrate that PYK specifically functions in the adult circadian clock to regulate period length.

Using the 35218 RNAi line showing the strongest phenotype, we confirmed specificity by overexpressing *Pyk* in the knockdown background, which restored both circadian period and *Pyk* mRNA levels (Fig. 1G and 1H). 35218 RNAi lines were used in the subsequent experiments.

To verify that adult-specific *Pyk* KD disrupts glycolysis, we performed metabolomic analysis of fly heads collected at ZT2 and ZT14 under 12:12LD cycle (Fig. 1I). *Pyk* KD flies exhibited elevated phosphoenolpyruvate (PEP) levels and reduced pyruvate (PYR) levels (substrate and product of PYK reaction, respectively), along with decreased NADH levels, confirming effective glycolytic impairment in circadian clock neurons.

### Adult-specific temporary KD of *Pyk* reduces PER and PDF levels

To investigate the mechanism underlying circadian period shortening in adult-specific *Pyk* KD flies, we examined clock protein levels by immunostaining for PER (transcriptional repressor in the TTFL) and PDF (neuropeptide in the output pathway) (Fig. 2). *Pyk* AKD flies showed reduced PER intensity in clock neurons, including LN_v_s, DN_1_s, and LN_d_s compared to controls at both ZT2 and ZT14 (Fig. 2A). Quantification of PER intensity in sLN_v_s across the circadian cycle confirmed significant reduction in *Pyk* AKD flies (Fig. 2B and 2C).

**Figure 2.**
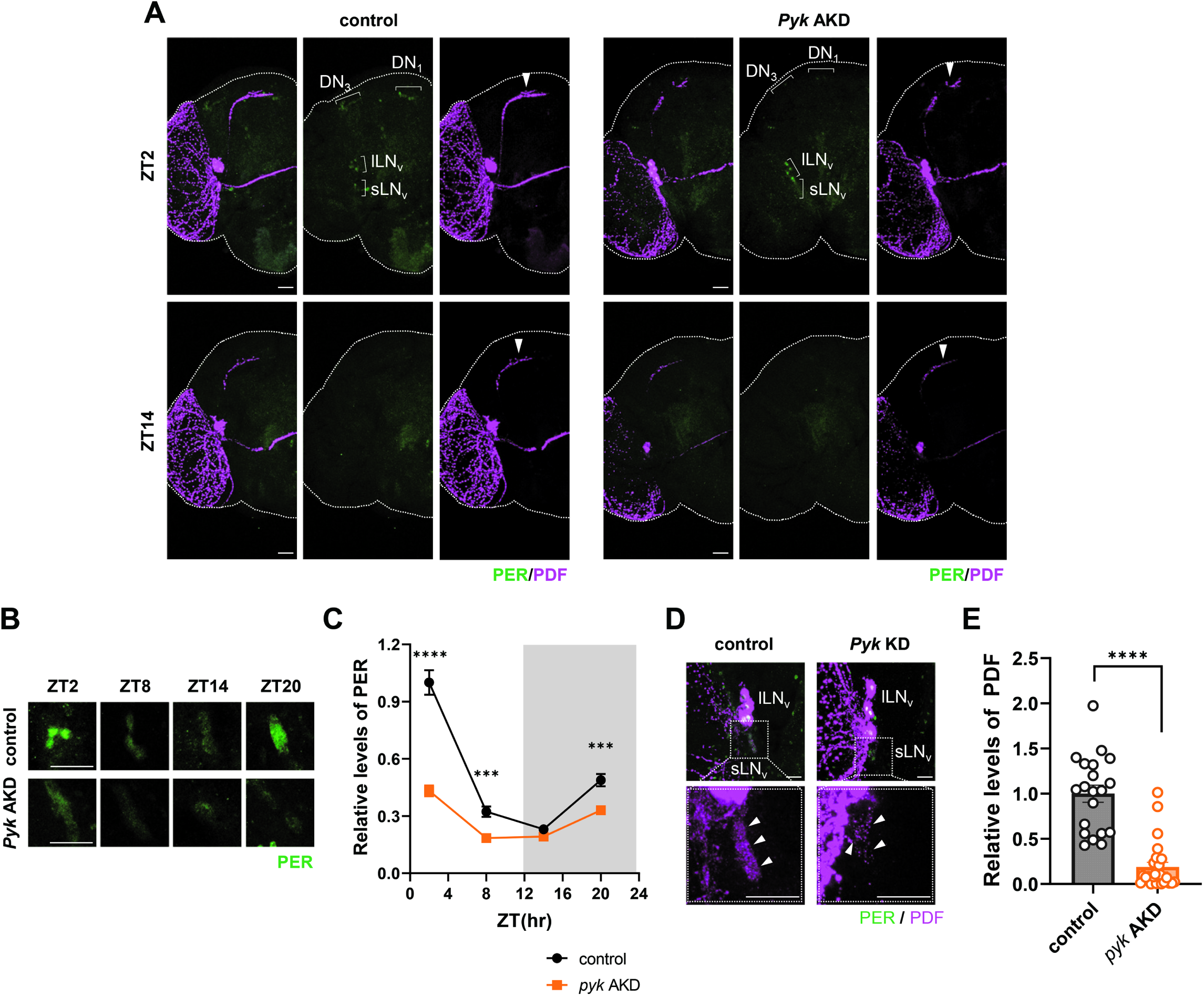
PER and PDF levels were reduced in *Pyk* AKD flies. (A – E) *dcr2*;tub-g80^ts^;TUG flies were crossed with *w*^1118^ (control) or UAS-*Pyk* Ri (*Pyk* AKD) flies and reared at 20°C. F1 offspring flies were maintained on a 12L:12D cycle at 30°C for 7 days and brains were dissected at ZT2 and ZT14 (A, C) or indicated times (B) and stained with anti-PER (Rb1, green) and anti-PDF (C7, magenta) antibodies. (A) The PER intensities in clock cells were reduced in *Pyk* AKD flies compared to control flies, and PDF intensities of sLN_v_ dorsally projecting neurites (arrow heads) were reduced in *Pyk* AKD flies compared to other flies. (B) Brains were dissected at indicated ZTs, and PER signal of sLN_v_s are shown. (C) PER intensities were quantified, and values indicate the mean ± SEM (n = 24–31). Asterisks indicate differences vs. control at each time points (two-way ANOVA with Sidak’s test): \*\*\**p* < 0.001, \*\*\*\**p* < 0.0001. (D) Brains were dissected at ZT2, and PER and PDF signal of LN_v_s are shown. Lower panels show a magnified image of the boxed region in the upper panel. The arrow heads indicate sLN_v_s. (E) PDF intensities were quantified, and bars indicate the mean ± SEM (n = 18–25). Statistical differences were determined using Mann-Whitney test: \*\*\*\**p* < 0.0001. All scale bars represent 20 *μ*m.

Additionally, *Pyk* AKD flies exhibited diminished PDF staining in sLN_v_ dorsal projections (Fig. 2A, arrowhead) and soma (Fig. 2D and 2E). Since reduced PER levels typically correlate with lengthened circadian periods (Bu *et al*, 2019; Cho *et al*., 2019; Lim *et al*, 2011; Richier *et al*, 2008; Xue *et al*, 2019), while reduced PDF is associated with shortened periods (Renn *et al*, 1999; Shafer & Taghert, 2009), we propose that the dominant effect of diminished PDF overrides the period-lengthening influence of reduced PER, resulting in net period shortening in *Pyk* AKD flies.

### PDF levels are highly responsive to *Pyk* expression

The profound decrease in PDF levels following adult-specific *Pyk* KD prompted investigation of whether this resulted from transient metabolic inhibition or permanent cellular damage. We conditionally knocked down *Pyk* for five days at 30°C, then restored expression by shifting flies to 20°C for five additional days. PDF levels, significantly reduced under *Pyk* AKD conditions, recovered to control levels following temperature shift (Fig. 3A, 3B), demonstrating that PDF levels dynamically respond to *Pyk* activity and that sLN_v_s retain functional integrity.

**Figure 3.**
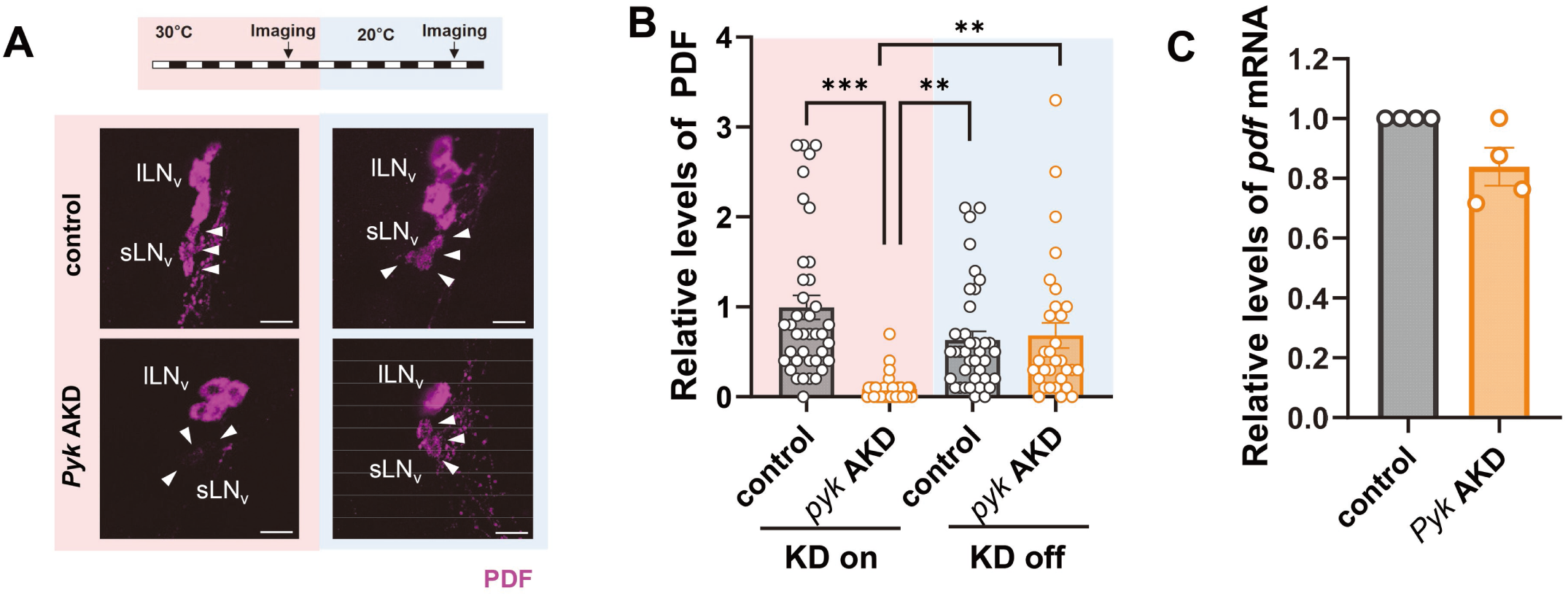
PDF levels were highly responsive to *Pyk* expression. (A - C) *dcr2*;tub-g80^ts^;TUG flies were crossed with *w*^1118^ (control) or UAS-*Pyk* Ri (*Pyk* AKD) flies and reared at 20°C. (A) F1 offspring were maintained on a 12L:12D cycle at 30°C (KD on, red background) for five days and then shifted to 20°C (KD off, blue background) for five days. Brains were dissected at ZT2 on day 5 (arrow, “Imaging”) and stained with anti-PDF (C7, magenta) antibodies. PDF intensities were significantly reduced in *Pyk* AKD flies compared to control at 30°C, but similar between control and *Pyk* AKD flies after temperature shift to 20°C. Arrowheads indicate sLN_v_s. All scale bars represent 20 *μ*m. (B) PDF intensities were quantified, and bars indicate the mean ± SEM (n = 31–39). Statistical differences were determined using one-way ANOVA with Tukey’s test: \*\**p* < 0.01, \*\*\*\**p* < 0.0001. (C) Flies were maintained on a 12L:12D cycle at 30°C for seven days and collected at ZT2. *Pdf* mRNA levels were quantified by real-time qRT-PCR and normalized to control. Bars indicate the mean ± SEM from four independent experiments. Statistical differences were determined using Mann–Whitney test.

To assess whether the reduction in PDF protein reflected transcriptional changes, we measured *Pdf* mRNA levels. *Pyk* AKD flies showed a modest, non-significant decrease in *Pdf* transcripts (Fig. 3C). Because the reduction in PDF protein was much larger than the change in mRNA, these data suggest that PYK regulates PDF primarily through post-transcriptional mechanisms.

### TARANIS levels are greatly reduced in *Pyk* KD flies

VRI functions as both a key TTFL component repressing *Clk* transcription and a regulator of circadian output supporting PDF expression (Cyran *et al*, 2003; Gunawardhana & Hardin, 2017). We initially investigated whether altered VRI expression could account for reduced PDF levels in *Pyk* KD flies. However, VRI levels in sLN_v_s at ZT14 - peak expression time - were similar between control and *Pyk* KD flies, indicating that VRI protein abundance is not directly correlated with PDF expression (Fig. 4A and 4B). A recent study found that *tara* mutants exhibited significantly reduced PDF levels (Akpoghiran *et al*., 2024). TARA, a homolog of the vertebrate TRIP-Br (Transcriptional Regulators Interacting with Plant homeodomain (PHD) zinc fingers and/or Bromodomains) proteins, contains multiple protein-interacting motifs and functions as a transcriptional co-regulator mediating interactions between transcription factors and chromatin remodeling proteins. It contributes to cell cycle regulation (Calgaro *et al*, 2002; Hsu *et al*, 2001) and circadian behavior (Akpoghiran *et al*., 2024), while its mammalian homolog TRIP-BR2 supports energy metabolism by regulating thermogenesis genes in adipocytes (Liew *et al*, 2013).

**Figure 4.**
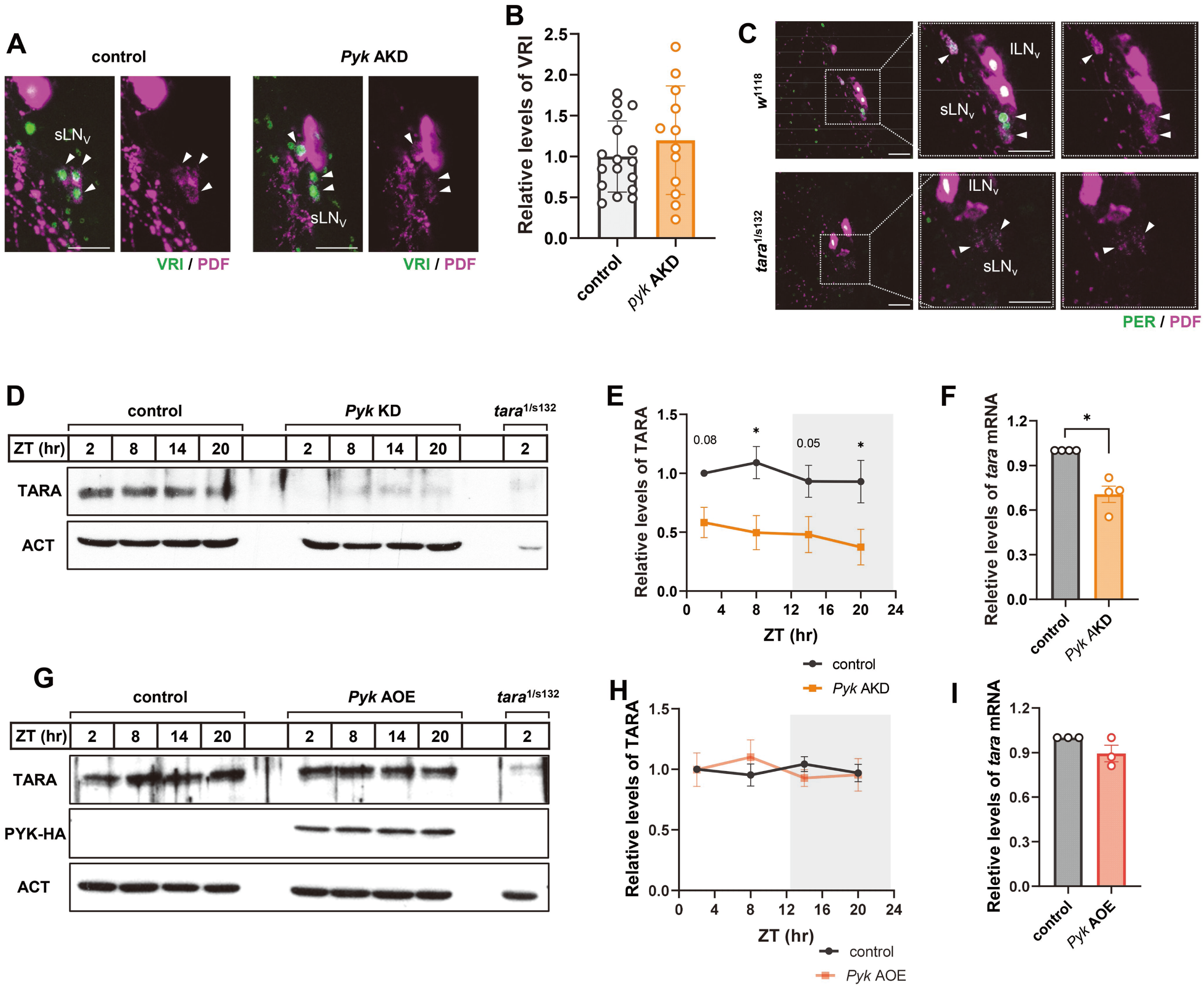
TARA levels were reduced in *Pyk* AKD flies. (A and B) *dcr2*;tub-g80^ts^;TUG flies were crossed with *w*^1118^ (control) or UAS-*Pyk* Ri (*Pyk* AKD) flies. F1 offsprings were maintained on a 12L:12D cycle at 30°C on 7 days, and brains were dissected at ZT14 and stained with anti-VRI (gp2, green) and anti-PDF (C7, magenta) antibodies. (A) The VRI expression in sLN_v_ of *Pyk* AKD was not different compared to control. (B) VRI intensities were quantified, and bars indicate the mean ± SEM (n = 12–17). (C) *w*^1118^ control and *tara*^1/s132^ hypomorphic mutant flies were maintained on a 12L:12D cycle at 25°C. Brains were dissected at ZT2 and stained with anti-PER (Rb1, green) and anti-PDF (C7, magenta) antibodies. The two right panels show magnified images of the boxed regions in the two left panels. Note that PDF intensities in sLN_v_ were greatly reduced in *tara*^1/s132^ flies. Arrowheads indicate sLN_v_s. All scale bars represent 20 *μ*m. (D - I) *dcr2*;tub-g80^ts^;TUG flies were crossed with *w*^1118^ (control) or UAS-*Pyk* Ri (*Pyk* AKD), or UAS-*Pyk*-HA (*Pyk* AOE) flies and reared at 20°C. (D, E, G, H) F1 offsprings and *tara*^1/s132^ were maintained on a 12L:12D cycle at 30°C for seven days, and protein extracts from the flies heads were processed for western blotting using anti-TARA antibodies. Actin served as the loading control. (E, H) TARA protein levels were quantified by measuring band intensities, and values indicate the mean ± SEM (n = 4). Asterisks indicate differences vs. control at each time points (two-way ANOVA with Sidak’s test): \**p* < 0.05. (F, I) Flies were maintained on a 12L:12D cycle at 30°C for seven days and collected at ZT2. *tara* mRNA levels were quantified by real-time qRT-PCR and normalized to control. Bars indicate the mean ± SEM from four independent experiments. Statistical differences were determined using Mann–Whitney test: \**p* < 0.05. Note that TARA protein and mRNA levels were greatly reduced in *Pyk* AKD flies but remained unchanged in *Pyk* AOE flies.

Given TARA’s roles in circadian behavior and metabolism, we hypothesized a potential link between TARA and PDF regulation via PYK. Consistent with previous reports (Akpoghiran *et al*., 2024), strongly hypomorphic *tara*^1/s132^ mutants (Afonso *et al*, 2015) showed markedly reduced PDF levels in sLN_v_s (Fig. 4C). Western blot analysis revealed that TARA protein levels, while constant across the circadian cycle as previously reported (Afonso *et al*., 2015), were substantially lower in *Pyk* AKD flies compared to controls (Fig. 4D – 4F). This reduction likely underestimates the extent of depletion in clock neurons, since TARA is expressed in all neurons (Afonso *et al*., 2015), while *Pyk* was specifically downregulated in *tim*-expressing cells. Additionally, *tara* mRNA levels were significantly reduced in *Pyk* AKD flies (Fig. 4F), indicating transcriptional regulation of TARA by PYK.

To test whether TARA expression responds to increased PYK levels, we overexpressed HA-tagged *Pyk* using the TUG;tub-gal80^ts^ driver (*Pyk* AOE). Both TARA protein (Fig. 4G and 4H) and mRNA (Fig. 4I) levels remained unchanged in *Pyk* AOE flies, suggesting that PYK was expressed at saturating levels under standard conditions. These findings demonstrate a crucial role for PYK in regulating TARA expression, which subsequently influences PDF levels.

### Expression of *tara* rescues the short period and restores PDF levels in P*yk* KD flies

To verify TARA’s role in regulating PDF levels in *Pyk* KD flies, we tested whether *tara* overexpression could rescue the short period phenotype. Adult-specific *tara* overexpression using TUG;tub-gal80^ts^, normalized the shortened circadian period caused by *Pyk* AKD, while *tara* overexpression alone did not affect period length (Fig. 5A). Although a previous study reported period lengthening with *tara* overexpression (Akpoghiran *et al*., 2024), that effect occurred at 25°C, whereas our experiments were conducted at 30°C. To assess temperature dependence, we overexpressed *tara* in LN_v_s at both temperatures. Results showed significant period lengthening at 25°C but not at 30°C (Fig. 5B), despite similar TARA::GFP fusion protein levels at both temperatures (Fig. 5C), indicating that the temperature sensitivity is not due to differences in protein abundance. Taken together, these data demonstrate that restoring TARA levels is sufficient to rescue *P*yk KD-induced period shortening.

**Figure 5.**
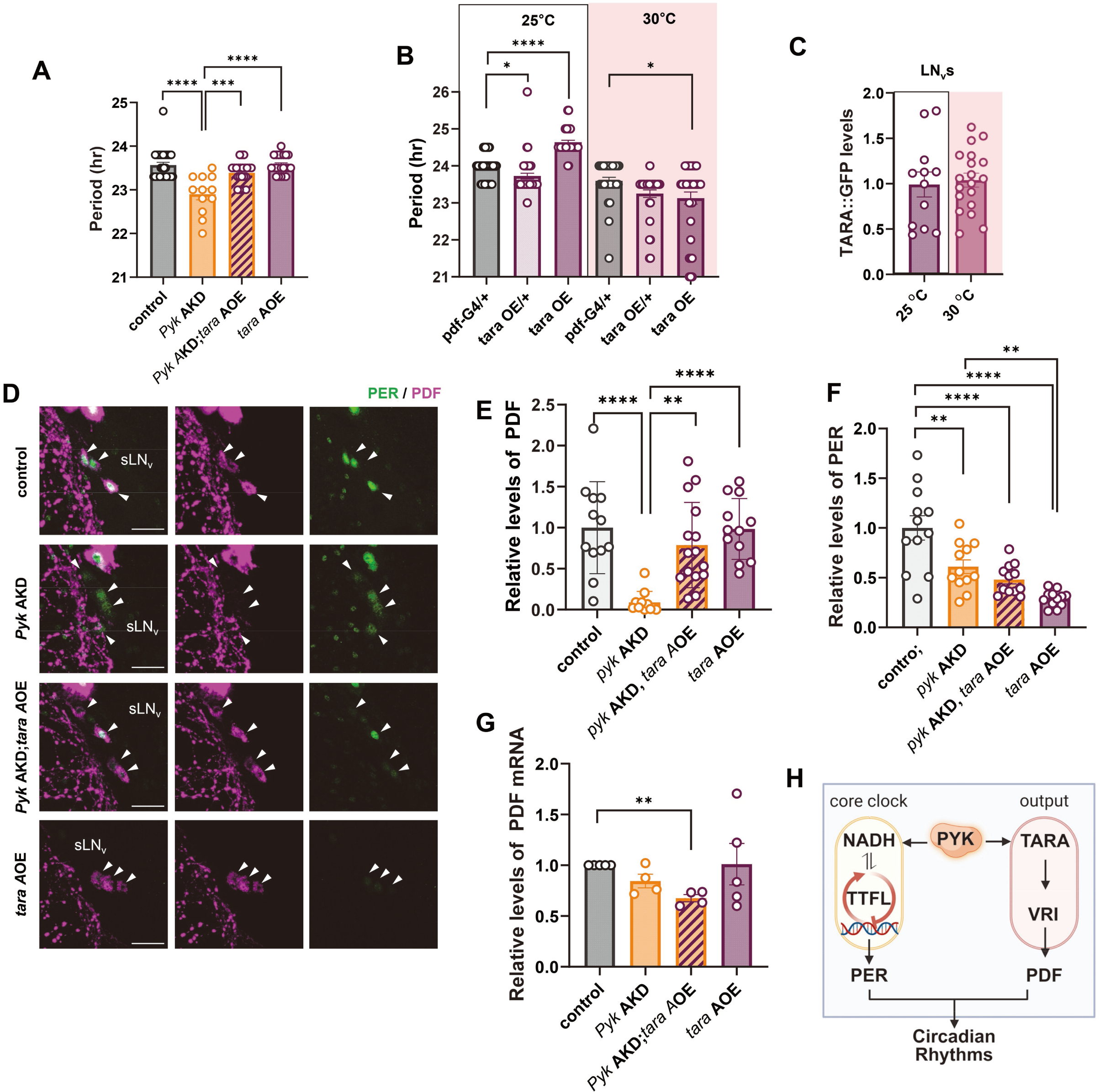
Overexpression of *tara* rescued the short-period phenotype and restored PDF levels in *Pyk* AKD flies. (A and B) *dcr2*;tub-g80^ts^;TUG flies were crossed with *w*^1118^ (control), UAS-*Pyk* Ri (*Pyk* AKD), or UAS-*tara* (*tara A*OE) either individually or in combination as indicated. Circadian rhythmic behavior of F1 offsprings were analyzed at 30°C. (A) Free-running periods of flies were shown. Bars indicate the mean ± SEM (n = 12–29). Statistical differences were determined using one-way ANOVA with Tukey’s test: \*\*\**p* < 0.001, \*\*\*\**p* < 0.0001. (B and C) *Pdf*-gal4 flies were crossed with *w*^1118^ (control) or UAS-*tara* (*tara* OE). (B) Circadian rhythmic behavior of F1 offsprings were analyzed at 25°C and 30°C. Free-running periods were shown. Bars indicate the mean ± SEM (n = 31–40). Statistical differences were determined using one-way ANOVA with Dunnet’s test: \**p* < 0.05, \*\*\*\**p* < 0.0001. (C) TARA levels in sLN_v_s were quantified by measuring GFP intensities which showed no significant differences between 25°C and 30°. Bars indicate the mean ± SEM (n = 12–19). (D - G) *dcr2*;tub-g80^ts^;TUG flies were crossed with *w*^1118^ (control), UAS-*Pyk* Ri (*Pyk* AKD), or UAS-*tara* (*tara* AOE) either individually or in combination as indicated. F1 offsprings were maintained under a 12L:12D cycle at 30°C. (D) Brains were dissected at ZT2 and stained with anti-PER (Rb1, green) and anti-PDF (C7, magenta) antibodies. Arrowheads indicate sLN_v_, and all scale bars represent 20 *μ*m. (E and F) PDF (E) and PER (F) intensities were quantified, and bars indicate the mean ± SEM (n = 11–15). Statistical differences were determined using one-way ANOVA with Dunnet’s test: \*\**p* < 0.01, \*\*\*\**p* < 0.0001. (G) Fly heads were collected at ZT2, and the *Pdf* mRNA levels were quantified using real-time qRT-PCR and normalized to the average level in control flies. Bars indicate mean ± SEM from four independent experiments. Asterisks indicate differences vs. control (*t* - test): \*\**p* < 0.001 (H) Schematic diagram illustrating our model of how PYK affects circadian rhythm. PYK knockdown reduces *tara* transcription, leading to decreased PDF levels via downregulation of VRI activity. Additionally, knockdown of PYK lowers NADH levels, resulting in reduced PER abundance. Created in https://BioRender.com

We next examined whether behavioral rescue correlated with restored PDF and PER levels in sLN_v_s. *tara* expression substantially restored reduced PDF levels in *Pyk* KD flies (Fig. 5D and 5E). Notably, *tara* overexpression alone did not increase PDF levels, suggesting that TARA and PYK function in the same pathway to restore PDF levels and are not additive (Fig. 5D and 5E). In contrast, PER levels were not rescued by TARA expression, supporting the idea that the period-shortening in *Pyk* KD flies results primarily from reduced PDF levels (Fig. 5D and 5F). Despite robust PDF protein restoration, *Pdf* mRNA remained low in *Pyk* KD flies with *tara* overexpression (Fig. 5G), suggesting that TARA enhances PDF expression post-transcriptionally in this context. This contrasts with previous findings that *tara* mutants exhibited reduced PDF at both transcriptional and post-transcriptional levels (Akpoghiran *et al*., 2024). TARA may normally exert both transcriptional and post-transcriptional control, but under hypometabolic conditions such as *Pyk* KD, its influence may shift toward post-transcriptional regulation.

In summary, our results demonstrate that PYK promotes *tara* expression, and that TARA overexpression is sufficient to restore PDF levels and rescue the period-shortening effects of *Pyk* KD, identifying TARA as a key mediator of glycolytic control over circadian output.

## Discussion

Circadian dysfunction and glucose hypometabolism are common in aging and neurodegenerative diseases, but the mechanistic connections between these processes have remain elusive. In this study, we demonstrate that temporarily knocking down key glycolytic enzyme *Pyk* in *Drosophila* clock cells shortened circadian period. Specifically, *Pyk* KD reduces levels of the clock protein PER as well as the output neuropeptide PDF. Mechanistically, these effects are separable; PDF reduction and circadian period shortening are mediated via TARA, while PER reduction appears to be TARA-independent and likely reflects a distinct, NADH-dependent mechanism (Fig. 5H).

Our findings establish TARA as a critical mediator of the effects of *Pyk* KD on PDF expression and circadian period. TARA modulates diverse cellular processes, including cell cycle (Calgaro *et al*., 2002; Hsu *et al*., 2001), sleep (Afonso *et al*., 2015), and circadian behavior (Akpoghiran *et al*., 2024). Recent studies showed that TARA modulates circadian period by enhancing the transcriptional activity of VRI and PDP1 (Akpoghiran *et al*., 2024). Additionally, VRI is essential for PDF expression at both the transcriptional and post-transcriptional levels and its adult-specific loss causes a shorter circadian period (Gunawardhana & Hardin, 2017). Integrating these findings, we propose that TARA reduction in *Pyk* KD flies leads to decreased VRI activity, resulting in reduced PDF levels and shortened circadian period (Fig. 5H). Several negative regulators of *Pdf* expression have been identified, including *Hormone receptor-like 38* as well as *stripe* and *scarecrow* (Mezan *et al*, 2016; Nair *et al*, 2020), and if they are upregulated by *Pyk* KD, it could contribute to lowered PDF levels. However, the restoration of PDF levels by *tara* expression in *Pyk* KD conditions demonstrates that TARA plays a central role in controlling PDF expression in response to PYK levels.

*Pyk* KD decreased *tara* mRNA levels, indicating that PYK regulates *tara* transcription. This extends the functional repertoire of glycolytic enzymes beyond metabolism. Beyond its canonical role in phosphorylating pyruvate, PYK can phosphorylate other protein substrates through direct interaction, altering their activity or stability (Akpoghiran *et al*., 2024; He *et al*, 2016; Zhao *et al*, 2018). However, co-immunoprecipitation assays failed to detect direct interactions between PYK and TARA, neither *in vitro* nor in flies, under our experimental conditions (data not shown), suggesting that PYK influences TARA levels through indirect mechanisms. PYK is known to perform non-glycolytic functions in the nucleus, where it influences transcription. The pyruvate kinase isoform of M2 (PKM2) translocates to the nucleus and interacts with various transcriptional regulators, promoting transcription of genes crucial for cell proliferation (Gao *et al*, 2012; Luo *et al*, 2011; Yang *et al*, 2012; Zhao *et al*., 2018). The reduction of *tara* mRNA in *Pyk* KD conditions supports the hypothesis that PYK influences TARA levels by regulating *tara* transcription (Fig. 4F). Given the critical role of TARA and its mammalian homologs, TRIP-Br proteins, in cell proliferation (Calgaro *et al*., 2002; Hsu *et al*., 2001), *tara* may represent a key transcriptional target of PYK.

Mammalian TARA homologs have been implicated in metabolic regulation. For example, TRIP-Br2 promotes lipid accumulation by enhancing the transcription of lipogenic genes (Liew *et al*., 2013), while TRIP-Br3 is downregulated under serum starvation to regulate apoptosis (Li *et al*, 2015). These roles support a model in which TARA and Trip-Br family proteins couple nutrient status to gene expression and cellular physiology. Our findings extend this model to circadian regulation, showing that TARA responds to changes in glucose metabolism and regulates neuropeptide expression, thereby linking energy state to behavioral rhythms. TARA and its mammalian homologs are known as transcriptional coregulators, consistent with a previous study showed TARA’s role in enhancing transcriptional activity of VRI and PDP1 (Akpoghiran *et al*., 2024). However, their functions are not limited to transcriptional coregulation. For example, TARA binds Cyclin A in the cytoplasm to regulate sleep (Afonso *et al*., 2015), and TRIP-Br1 binds XIAP to ubiquitinate and degrade Adenylyl Cyclases (Hu *et al*, 2017). Thus, TARA and TRIP-Br proteins may function as a context-dependent integrator of metabolic and circadian signals, with both nuclear and cytoplasmic roles.

Metabolmic analyses confirmed that temporary *Pyk* KD in clock cells induces hypometabolic state, characterized by reduced pyruvate and NADH levels, (Fig. 1I). This hypometabolic state greatly reduced PER levels (Fig. 2B). While TARA overexpression was sufficient to reverse the shorter period and reduced PDF expression by *Pyk* KD, it was unable to restore PER expression, suggesting that PYK impacts PER expression through a TARA-independent pathway. The relationship between cellular redox state and circadian clock function is well-established in mammals, where NADH levels correlate with circadian clock activity by enhancing CLOCK:BMAL1 DNA binding activity (Rutter *et al*., 2001). The NAD+ dependent histone deacetylase sirtuin1 (SIRT1) interacts with CLOCK:BMAL1 and antagonizes CLOCK’s histone acetyltransferase (HAT) activity, thereby inhibiting CLOCK:BMAL1 function (Nakahata *et al*., 2008). SIRT1 also deacetylates PER, reducing its stability (Asher *et al*., 2008; Nakahata *et al*, 2009). Additionally, oxidized cellular redox states following prolonged pentose phosphate pathway inhibition dramatically reduce per2::luc rhythm amplitudes in mouse fibroblasts (Putker *et al*, 2018). Given the highly conserved nature of circadian clock mechanism, the observed decrease in NADH levels following *Pyk* KD in flies likely contributes to reduced PER levels through similar mechanisms (Fig. 5H). Notably, ATP levels remained unaffected in *Pyk* KD conditions, suggesting that ATP production may be maintained through alternative metabolic pathways, such as fatty acid oxidation, which do not require pyruvate.

The clinical relevance of our findings is underscored by the observation that regional glucose deficits in the brain occur with aging and precede neurodegenerative diseases, such as Alzheimer’s disease and Parkinson’s disease (Lewitt *et al*, 2013; Mosconi *et al*, 2006; Peppard *et al*, 1992; Small *et al*, 2000; Xu *et al*, 2015). Circadian period changes with aging have been variably reported-with some studies noting no change (Czeisler *et al*, 1999; Davis & Viswanathan, 1998; Sharma & Chandrashekaran, 1998) or lengthening (Kendall *et al*, 2001; Possidente *et al*, 1995; Valentinuzzi *et al*, 1997)- a number of reports support a trend toward period shortening (Pittendrigh & Daan, 1974; van Gool *et al*, 1987; Weitzman *et al*, 1982; Zee *et al*, 1992). Since the expression of glycolytic enzymes, including PYK, is regulated by glucose availability (Greiner *et al*, 1994; Marie *et al*, 1993; Roche *et al*, 1997), reduced glucose uptake in aging and neurodegenerative diseases might decrease glycolytic enzyme expression, paralleling our temporary *Pyk* KD condition. Importantly, circadian clock gene expression becomes dysregulated in aging and neurodegenerative diseases(Bonaconsa *et al*, 2014; Chen *et al*, 2016; Kolker *et al*, 2003; Nakamura *et al*, 2015; Rakshit *et al*, 2012; Stevanovic *et al*, 2017; Wang *et al*, 2016). Moreover, VIP signaling is reduced in the mammalian SCN under these conditions (Arai *et al*, 1984; Chee *et al*, 1988; Fahrenkrug *et al*, 2007; Roozendaal *et al*, 1987; Stopa *et al*, 1999; van Wamelen *et al*, 2013; Zhou *et al*, 1995). Similarly, aged *Drosophila* show reduced PDF expression (Umezaki *et al*, 2012). Our study clearly demonstrates that glucose hypometabolism in clock cells reduces both PER and PDF levels, providing mechanistic support for these clinical observations.

In summary, our study enhances understanding of the circadian rhythm disruptions by glucose hypometabolism by establishing the PYK-TARA-PDF regulatory axis. By identifying this regulatory pathway, we show that nutrient status can shape circadian rhythms through multiple molecular mechanisms. Circadian disruption in aging and disease may reflect the combined effects of redox imbalance and impaired neuropeptide signaling due to metabolic dysfunction.

## Materials and Methods

### Fly stocks

*tara*^1^, *tara*^S132^ and UAS-*tara* flies have been previously reported (Afonso *et al*., 2015; Akpoghiran *et al*., 2024). The *Pyk*-HA (Ham *et al*, 2021) flies were a gift from Jongkyeong Chung (Seoul National University, Republic of Korea). The following fly lines were obtained from the Bloomington Drosophila Stock Center: *w*^1118^(BL5905), *Hex*-A RNAi (BL35155), *Hex*-C RNAi (BL57404, BL35338, BL35337), *Pfk* RNAi (BL34336, BL36782), *Pyk* RNAi (BL35218) and *Pdf*-gal4 (BL80939). The following fly lines were obtained from the Vienna Drosophila Resource Center: *Hex*-A RNAi (21054, 104680), *Pfk* RNAi (105666), *Pyk* RNAi (35165, 49533). *tim*(UAS)-gal4 (TUG), and TUG;tub-gal80^ts^ were crossed to UAS-*dicer2 (dcr2)*/CyO to generate *dcr2*;TUG (*d2*;TUG) and *dcr2*;tub-g80^ts^;TUG, which were used for knockdown of expression.

### Locomotor behavior analysis

Locomotor activity of individual flies was determined using the *Drosophila* Activity Monitoring System (Trikinetics). 2 - 5 days old young male flies were used for the analysis and maintained in glass tubes containing 2% agar and 5% sucrose. Flies were kept in incubators at the indicated temperature (20°C, 25°C or 30°C) under 12L:12D cycle for 4 days and then were maintained under constant dark (DD) condition for 7 days. Circadian rhythmic behavior was analyzed using FaasX software (Fly Activity Analysis Suite for MacOSX), which was generously provided by Francois Rouyer (Centre National de la Recherche Scientifique, France). Periods were calculated for each fly using χ^2^ periodogram analysis, and data were pooled to obtain an average value. The power was calculated by quantifying the relative strength of the rhythm during the DD condition. Individual flies with a power ≥10 and width ≥2 were considered rhythmic.

### Metabolomics analysis

Fly heads were homogenized in methanol using a TissueLyzer (Qiagen). Internal standard solution (malonyl-^13^C_3_ CoA, 5 μM Gln-d_4_) was added to the samples. Following centrifugation at 13000 rpm for 10 minutes (Eppendorf Centrifuge 5415R), the resulting precipitate was reserved for later protein quantification. The supernatant was then subjected to liquid-liquid extraction, and the aqueous phase was collected for metabolomic analysis. Metabolites were analyzed using LC-MS/MS (1290 HPLC (Agilent) coupled with Qtrap 5500 (ABSciex). For metabolites related to energy metabolism, the Synergi Fusion RP 50 × 2 mm column was used, with mobile phase A being 5 mM CH_3_COONH_4_ in H_2_O and mobile phase B being 5 mM CH_3_COONH_4_ in MeOH. The separation gradient was: 0% B for 5 min, 0% to 90% B for 2 min, hold at 90% for 8 min, 90% to 0% B for 1 min, and then hold at 0% B for 9 min. The flow rate was 70 *μl*/min, except between minutes 7 and 15, where it increased to 140 *μl*/min. The column temperature was maintained at 23 °C. Multiple reaction monitoring (MRM) was employed for analysis. The quantitative value of each metabolite was normalized to the total protein amount. To obtain relative levels of metabolites among samples, the metabolite level of control flies at ZT2 was set to 1 and other values were normalized.

### Immunohistochemistry and confocal imaging

Immunostaining was performed as described previously with minor modifications (Lee *et al*, 2016). Fly heads were cut open, fixed in 2% formaldehyde, and washed with 0.03% PAXD buffer (1X PBS, 5% BSA, 0.03% sodium deoxycholate, 0.03% Triton X-100) (Gunawardhana & Hardin, 2017). The fixed heads were dissected, and the isolated brains were permeabilized in 1% PBST for 20 min and then blocked in PAXD containing 5% horse serum for 1 hr. The following primary antibodies were diluted 1:200 and added directly to the mixtures:, anti-PER antibody (Rb1) (Kim *et al*, 2012), anti-PDF antibody (C7) (DSHB) and anti-VRI antibody (gp2) (Glossop *et al*, 2003). The brains were washed with PAXD and incubated overnight with secondary antibodies in a blocking solution at 4 °C. The following secondary antibodies were used at a 1:200 dilution: goat anti-rabbit Alexa-488 (Thermo Fisher Scientific) and goat anti-mouse Alexa-555 (Thermo Fisher Scientific). Stained brain samples were washed with PAXD, incubated in 0.1 M phosphate buffer containing 50% glycerol for 30 min, and mounted using a mounting medium. Confocal images were acquired using an LSM 800 confocal microscope (Carl Zeiss) and processed with Zen software (ZEN Digital Imaging for Light Microscopy, Carl Zeiss). The displayed representative images are Z-stack projections to improve visualization. For intensity quantification, single plane images were used.

### Western blotting

For Western blotting, protein extracts were prepared in RIPA lysis buffer [25 mM Tris-HCl, pH 7.5, 50 mM NaCl, 0.5% sodium deoxycholate, 0.5% NP-40, and 0.1% SDS] with freshly added protease inhibitor mixture (Sigma-Aldrich) and phosphatase inhibitor mixture (Sigma-Aldrich). Protein extracts were resolved by SDS-PAGE with the 10% polyacrylamide gel. Primary antibodies were used at the following dilutions: anti-TARA (rat), 1:3000 (Afonso *et al*., 2015), anti-actin (Rb) (Sigma-Aldrich), 1:3000 and anti-HA (Rat) (Roche Diagnostics), 1:3000. Band intensity was quantified using ImageJ software.

### qRT-PCR

Total RNA was extracted from fly heads using QIAzol reagent (QIAGEN). Total RNA (1 μg) was reverse transcribed using an oligo(dT)20 primer (for mRNA) and PrimeScript RTase (TaKaRa). Quantitative, real-time PCR (qPCR) was performed using Rotor Gene 6000 (QIAGEN) with TB Green Premix Ex Taq (Tli RNaseH Plus, TaKaRa). The following primers were used: *hex* forward, 5-CGAGAACTTATGCAACCCT-3; *hex* reverse, 5-AGCGACTGTACACTTCCTG-3; *pfk* forward, 5-GGCAAGCCCAAAACGGAAAT-3; *pfk* reverse, 5-CGACATAGTACTGCGGCCAT-3; *pyk* forward, 5-CTCATCTACAAGGAGCCCG-3; *pyk* reverse, 5-TCTTCTTTCCGACCTGCAG-3; *tara* forward, 5-GTGCACTGAGGTGAATTCC-3; *tara* reverse, 5-ATCCTTGCTGTCGAAGGTC-3; *Pdf* forward, 5-GCCACTCTCTGTCGCTATCC-3; and *Pdf* reverse, 5-CAGTGGTGGGTCGTCCTAAT-3. Noncycling mRNA encoding *cbp20* was used to normalize gene expression with the primers *cbp20* forward, 5-GTATAAGAAGACGCCCTGC-3; and *cbp20* reverse, 5-TTCACAAATCTCATGGCCG-3. The data were analyzed using Rotor Gene Q - Pure Detection software (2.2.3), and the relative mRNA levels were quantified using the 2^-ΔΔCt^ method in which ΔΔCt = [(C_t_ target - C_t_ *cbp20*) of experimental group] - [(C_t_ target - C_t_ *cbp20*) of control group].

### Statistics and Reproducibility

Statistical analysis was conducted using GraphPad Prism 8 software. To assess data distribution, the Shapiro-Wilk normality test was applied (*p* < 0.05). One-way ANOVA with Dunnett’s test or Tukey’s test as a post-hoc analysis was performed in multiple groups. When the number of samples is too small (less than 5), to enhance the statistical powers and sensitivity, comparisons between experimental groups were made using either the Student’s t-test for data for normally distributed data or the Mann-Whitney U test for non-normally distributed data.

## Acknowledgements

We are very grateful to Jongkyeong Chung (Seoul National University, Republic of Korea) for providing UAS-*Pyk*-HA flies and Paul E. Hardin (Texas A&M University, USA) for providing anti-VRI antibodies. We also thank the members of the KEY laboratory for their insightful discussions and comments. *Drosophila* stocks obtained from the Bloomington Drosophila Stock Center (NIH P40OD018537) and Vienna Drosophila Resource Center (VDRC, www.vdrc.at). were used in this study. This research was funded by grants from the National Research Foundation of Korea, M-2024-A0403-00020 (to S.H.L) and NRF-2019R1A5A2026045, RS-2023-00208490 (to E.Y.K), as well as the National Institutes of Health of USA (R01NS086887 to K.K.)

## References

Afonso DJ, Liu D, Machado DR, Pan H, Jepson JE, Rogulja D, Koh K (2015) TARANIS Functions with Cyclin A and Cdk1 in a Novel Arousal Center to Control Sleep in Drosophila. Curr Biol 25: 1717–1726

Akpoghiran O, Afonso DJS, Zhang Y, Koh K (2024) TARANIS Interacts with VRILLE and PDP1 to Modulate the Circadian Transcriptional Feedback Mechanism in Drosophila. J Neurosci 44

Arai H, Moroji T, Kosaka K (1984) Somatostatin and vasoactive intestinal polypeptide in postmortem brains from patients with Alzheimer-type dementia. Neurosci Lett 52: 73–78

Asher G, Gatfield D, Stratmann M, Reinke H, Dibner C, Kreppel F, Mostoslavsky R, Alt FW, Schibler U (2008) SIRT1 regulates circadian clock gene expression through PER2 deacetylation. Cell 134: 317–328

Bonaconsa M, Malpeli G, Montaruli A, Carandente F, Grassi-Zucconi G, Bentivoglio M (2014) Differential modulation of clock gene expression in the suprachiasmatic nucleus, liver and heart of aged mice. Exp Gerontol 55: 70–79

Bu B, He W, Song L, Zhang L (2019) Nuclear Envelope Protein MAN1 Regulates the Drosophila Circadian Clock via Period. Neurosci Bull 35: 969–978

Calgaro S, Boube M, Cribbs DL, Bourbon HM (2002) The Drosophila gene taranis encodes a novel trithorax group member potentially linked to the cell cycle regulatory apparatus. Genetics 160: 547–560

Camandola S, Mattson MP (2017) Brain metabolism in health, aging, and neurodegeneration. EMBO J 36: 1474–1492

Chee CA, Roozendaal B, Swaab DF, Goudsmit E, Mirmiran M (1988) Vasoactive intestinal polypeptide neuron changes in the senile rat suprachiasmatic nucleus. Neurobiol Aging 9: 307–312

Chen CY, Logan RW, Ma T, Lewis DA, Tseng GC, Sibille E, McClung CA (2016) Effects of aging on circadian patterns of gene expression in the human prefrontal cortex. Proc Natl Acad Sci U S A 113: 206–211

Cho E, Kwon M, Jung J, Kang DH, Jin S, Choi SE, Kang Y, Kim EY (2019) AMP-Activated Protein Kinase Regulates Circadian Rhythm by Affecting CLOCK in Drosophila. J Neurosci 39: 3537–3550

Cyran SA, Buchsbaum AM, Reddy KL, Lin MC, Glossop NR, Hardin PE, Young MW, Storti RV, Blau J (2003) vrille, Pdp1, and dClock form a second feedback loop in the Drosophila circadian clock. Cell 112: 329–341

Czeisler CA, Duffy JF, Shanahan TL, Brown EN, Mitchell JF, Rimmer DW, Ronda JM, Silva EJ, Allan JS, Emens JS et al (1999) Stability, precision, and near-24-hour period of the human circadian pacemaker. Science 284: 2177–2181

Davis FC, Viswanathan N (1998) Stability of circadian timing with age in Syrian hamsters. Am J Physiol 275: R960–968

Duvernell DD, Eanes WF (2000) Contrasting molecular population genetics of four hexokinases in Drosophila melanogaster, D. simulans and D. yakuba. Genetics 156: 1191–1201

Fahrenkrug J, Popovic N, Georg B, Brundin P, Hannibal J (2007) Decreased VIP and VPAC2 receptor expression in the biological clock of the R6/2 Huntington’s disease mouse. J Mol Neurosci 31: 139–148

Gao X, Wang H, Yang JJ, Liu X, Liu ZR (2012) Pyruvate kinase M2 regulates gene transcription by acting as a protein kinase. Mol Cell 45: 598–609

Glossop NR, Houl JH, Zheng H, Ng FS, Dudek SM, Hardin PE (2003) VRILLE feeds back to control circadian transcription of Clock in the Drosophila circadian oscillator. Neuron 37: 249–261

Gonzalez-Redondo R, Garcia-Garcia D, Clavero P, Gasca-Salas C, Garcia-Eulate R, Zubieta JL, Arbizu J, Obeso JA, Rodriguez-Oroz MC (2014) Grey matter hypometabolism and atrophy in Parkinson’s disease with cognitive impairment: a two-step process. Brain 137: 2356–2367

Greiner EF, Guppy M, Brand K (1994) Glucose is essential for proliferation and the glycolytic enzyme induction that provokes a transition to glycolytic energy production. J Biol Chem 269: 31484–31490

Gunawardhana KL, Hardin PE (2017) VRILLE Controls PDF Neuropeptide Accumulation and Arborization Rhythms in Small Ventrolateral Neurons to Drive Rhythmic Behavior in Drosophila. Curr Biol 27: 3442–3453 e3444

Ham SJ, Lee D, Xu WJ, Cho E, Choi S, Min S, Park S, Chung J (2021) Loss of UCHL1 rescues the defects related to Parkinson’s disease by suppressing glycolysis. Sci Adv 7

He CL, Bian YY, Xue Y, Liu ZX, Zhou KQ, Yao CF, Lin Y, Zou HF, Luo FX, Qu YY et al (2016) Pyruvate Kinase M2 Activates mTORC1 by Phosphorylating AKT1S1. Sci Rep 6: 21524

Herzog ED (2007) Neurons and networks in daily rhythms. Nat Rev Neurosci 8: 790–802

Holth JK, Fritschi SK, Wang C, Pedersen NP, Cirrito JR, Mahan TE, Finn MB, Manis M, Geerling JC, Fuller PM et al (2019) The sleep-wake cycle regulates brain interstitial fluid tau in mice and CSF tau in humans. Science 363: 880–884

Hood S, Amir S (2017a) The aging clock: circadian rhythms and later life. J Clin Invest 127: 437–446

Hood S, Amir S (2017b) Neurodegeneration and the Circadian Clock. Front Aging Neurosci 9: 170

Hsu SI, Yang CM, Sim KG, Hentschel DM, O’Leary E, Bonventre JV (2001) TRIP-Br: a novel family of PHD zinc finger- and bromodomain-interacting proteins that regulate the transcriptional activity of E2F-1/DP-1. EMBO J 20: 2273–2285

Hu W, Yu X, Liu Z, Sun Y, Chen X, Yang X, Li X, Lam WK, Duan Y, Cao X et al (2017) The complex of TRIP-Br1 and XIAP ubiquitinates and degrades multiple adenylyl cyclase isoforms. Elife 6

Kadener S, Stoleru D, McDonald M, Nawathean P, Rosbash M (2007) Clockwork Orange is a transcriptional repressor and a new Drosophila circadian pacemaker component. Genes Dev 21: 1675–1686

Kang JE, Lim MM, Bateman RJ, Lee JJ, Smyth LP, Cirrito JR, Fujiki N, Nishino S, Holtzman DM (2009) Amyloid-beta dynamics are regulated by orexin and the sleep-wake cycle. Science 326: 1005–1007

Kendall AR, Lewy AJ, Sack RL (2001) Effects of aging on the intrinsic circadian period of totally blind humans. J Biol Rhythms 16: 87–95

Kim EY, Jeong EH, Park S, Jeong HJ, Edery I, Cho JW (2012) A role for O-GlcNAcylation in setting circadian clock speed. Genes Dev 26: 490–502

King AN, Sehgal A (2020) Molecular and circuit mechanisms mediating circadian clock output in the Drosophila brain. Eur J Neurosci 51: 268–281

Kolker DE, Fukuyama H, Huang DS, Takahashi JS, Horton TH, Turek FW (2003) Aging alters circadian and light-induced expression of clock genes in golden hamsters. J Biol Rhythms 18: 159–169

Kress GJ, Liao F, Dimitry J, Cedeno MR, FitzGerald GA, Holtzman DM, Musiek ES (2018) Regulation of amyloid-beta dynamics and pathology by the circadian clock. J Exp Med 215: 1059–1068

Lamia KA, Sachdeva UM, DiTacchio L, Williams EC, Alvarez JG, Egan DF, Vasquez DS, Juguilon H, Panda S, Shaw RJ et al (2009) AMPK regulates the circadian clock by cryptochrome phosphorylation and degradation. Science 326: 437–440

Lananna BV, Musiek ES (2020) The wrinkling of time: Aging, inflammation, oxidative stress, and the circadian clock in neurodegeneration. Neurobiol Dis 139: 104832

Lee E, Cho E, Kang DH, Jeong EH, Chen Z, Yoo SH, Kim EY (2016) Pacemaker-neuron-dependent disturbance of the molecular clockwork by a Drosophila CLOCK mutant homologous to the mouse Clock mutation. Proc Natl Acad Sci U S A 113: E4904–4913

Lee SH, Cho E, Yoon SE, Kim Y, Kim EY (2021) Metabolic control of daily locomotor activity mediated by tachykinin in Drosophila. Commun Biol 4: 693

Leng Y, Musiek ES, Hu K, Cappuccio FP, Yaffe K (2019) Association between circadian rhythms and neurodegenerative diseases. Lancet Neurol 18: 307–318

Lewitt PA, Li J, Lu M, Beach TG, Adler CH, Guo L, Arizona Parkinson’s Disease C (2013) 3-hydroxykynurenine and other Parkinson’s disease biomarkers discovered by metabolomic analysis. Mov Disord 28: 1653–1660

Li C, Jung S, Lee S, Jeong D, Yang Y, Kim KI, Lim JS, Cheon CI, Kim C, Kang YS et al (2015) Nutrient/serum starvation derived TRIP-Br3 down-regulation accelerates apoptosis by destabilizing XIAP. Oncotarget 6: 7522–7535

Liew CW, Boucher J, Cheong JK, Vernochet C, Koh HJ, Mallol C, Townsend K, Langin D, Kawamori D, Hu J et al (2013) Ablation of TRIP-Br2, a regulator of fat lipolysis, thermogenesis and oxidative metabolism, prevents diet-induced obesity and insulin resistance. Nat Med 19: 217–226

Lim C, Chung BY, Pitman JL, McGill JJ, Pradhan S, Lee J, Keegan KP, Choe J, Allada R (2007) Clockwork orange encodes a transcriptional repressor important for circadian-clock amplitude in Drosophila. Curr Biol 17: 1082–1089

Lim C, Lee J, Choi C, Kilman VL, Kim J, Park SM, Jang SK, Allada R, Choe J (2011) The novel gene twenty-four defines a critical translational step in the Drosophila clock. Nature 470: 399–403

Linford NJ, Chan TP, Pletcher SD (2012) Re-patterning sleep architecture in Drosophila through gustatory perception and nutritional quality. PLoS Genet 8: e1002668

Liu JA, Bumgarner JR, Walker WH, 2nd, Melendez-Fernandez OH, Walton JC, DeVries AC, Nelson RJ (2024) Chronic phase advances reduces recognition memory and increases vascular cognitive dementia-like impairments in aged mice. Sci Rep 14: 7760

Luo W, Hu H, Chang R, Zhong J, Knabel M, O’Meally R, Cole RN, Pandey A, Semenza GL (2011) Pyruvate kinase M2 is a PHD3-stimulated coactivator for hypoxia-inducible factor 1. Cell 145: 732–744

Majcin Dorcikova M, Duret LC, Pottie E, Nagoshi E (2023) Circadian clock disruption promotes the degeneration of dopaminergic neurons in male Drosophila. Nat Commun 14: 5908

Marcheva B, Ramsey KM, Peek CB, Affinati A, Maury E, Bass J (2013) Circadian clocks and metabolism. Handb Exp Pharmacol: 127–155

Marie S, Diaz-Guerra MJ, Miquerol L, Kahn A, Iynedjian PB (1993) The pyruvate kinase gene as a model for studies of glucose-dependent regulation of gene expression in the endocrine pancreatic beta-cell type. J Biol Chem 268: 23881–23890

Matsumoto A, Ukai-Tadenuma M, Yamada RG, Houl J, Uno KD, Kasukawa T, Dauwalder B, Itoh TQ, Takahashi K, Ueda R et al (2007) A functional genomics strategy reveals clockwork orange as a transcriptional regulator in the Drosophila circadian clock. Genes Dev 21: 1687–1700

McGuire SE, Mao Z, Davis RL (2004) Spatiotemporal gene expression targeting with the TARGET and gene-switch systems in Drosophila. Sci STKE 2004: pl6

Mezan S, Feuz JD, Deplancke B, Kadener S (2016) PDF Signaling Is an Integral Part of the Drosophila Circadian Molecular Oscillator. Cell Rep 17: 708–719

Mohawk JA, Green CB, Takahashi JS (2012) Central and peripheral circadian clocks in mammals. Annu Rev Neurosci 35: 445–462

Mosconi L, Pupi A, De Leon MJ (2008) Brain glucose hypometabolism and oxidative stress in preclinical Alzheimer’s disease. Ann N Y Acad Sci 1147: 180–195

Mosconi L, Sorbi S, de Leon MJ, Li Y, Nacmias B, Myoung PS, Tsui W, Ginestroni A, Bessi V, Fayyazz M et al (2006) Hypometabolism exceeds atrophy in presymptomatic early-onset familial Alzheimer’s disease. J Nucl Med 47: 1778–1786

Moser D, Johnson L, Lee CY (1980) Multiple forms of Drosophila hexokinase. Purification, biochemical and immunological characterization. J Biol Chem 255: 4673–4679

Nair S, Bahn JH, Lee G, Yoo S, Park JH (2020) A Homeobox Transcription Factor Scarecrow (SCRO) Negatively Regulates Pdf Neuropeptide Expression through Binding an Identified cis-Acting Element in Drosophila melanogaster. Mol Neurobiol 57: 2115–2130

Nakahata Y, Kaluzova M, Grimaldi B, Sahar S, Hirayama J, Chen D, Guarente LP, Sassone-Corsi P (2008) The NAD+-dependent deacetylase SIRT1 modulates CLOCK-mediated chromatin remodeling and circadian control. Cell 134: 329–340

Nakahata Y, Sahar S, Astarita G, Kaluzova M, Sassone-Corsi P (2009) Circadian control of the NAD+ salvage pathway by CLOCK-SIRT1. Science 324: 654–657

Nakamura TJ, Nakamura W, Tokuda IT, Ishikawa T, Kudo T, Colwell CS, Block GD (2015) Age-Related Changes in the Circadian System Unmasked by Constant Conditions. eNeuro 2

Patke A, Young MW, Axelrod S (2020) Molecular mechanisms and physiological importance of circadian rhythms. Nat Rev Mol Cell Biol 21: 67–84

Peppard RF, Martin WR, Carr GD, Grochowski E, Schulzer M, Guttman M, McGeer PL, Phillips AG, Tsui JK, Calne DB (1992) Cerebral glucose metabolism in Parkinson’s disease with and without dementia. Arch Neurol 49: 1262–1268

Pittendrigh CS, Daan S (1974) Circadian oscillations in rodents: a systematic increase of their frequency with age. Science 186: 548–550

Possidente B, McEldowney S, Pabon A (1995) Aging lengthens circadian period for wheel-running activity in C57BL mice. Physiol Behav 57: 575–579

Putker M, Crosby P, Feeney KA, Hoyle NP, Costa ASH, Gaude E, Frezza C, O’Neill JS (2018) Mammalian Circadian Period, But Not Phase and Amplitude, Is Robust Against Redox and Metabolic Perturbations. Antioxid Redox Signal 28: 507–520

Rakshit K, Krishnan N, Guzik EM, Pyza E, Giebultowicz JM (2012) Effects of aging on the molecular circadian oscillations in Drosophila. Chronobiol Int 29: 5–14

Renn SC, Park JH, Rosbash M, Hall JC, Taghert PH (1999) A pdf neuropeptide gene mutation and ablation of PDF neurons each cause severe abnormalities of behavioral circadian rhythms in Drosophila. Cell 99: 791–802

Richier B, Michard-Vanhee C, Lamouroux A, Papin C, Rouyer F (2008) The clockwork orange Drosophila protein functions as both an activator and a repressor of clock gene expression. J Biol Rhythms 23: 103–116

Roche E, Assimacopoulos-Jeannet F, Witters LA, Perruchoud B, Yaney G, Corkey B, Asfari M, Prentki M (1997) Induction by glucose of genes coding for glycolytic enzymes in a pancreatic beta-cell line (INS-1). J Biol Chem 272: 3091–3098

Roozendaal B, van Gool WA, Swaab DF, Hoogendijk JE, Mirmiran M (1987) Changes in vasopressin cells of the rat suprachiasmatic nucleus with aging. Brain Res 409: 259–264

Rutter J, Reick M, Wu LC, McKnight SL (2001) Regulation of clock and NPAS2 DNA binding by the redox state of NAD cofactors. Science 293: 510–514

Schibler U, Gotic I, Saini C, Gos P, Curie T, Emmenegger Y, Sinturel F, Gosselin P, Gerber A, Fleury-Olela F et al (2015) Clock-Talk: Interactions between Central and Peripheral Circadian Oscillators in Mammals. Cold Spring Harb Symp Quant Biol 80: 223–232

Shafer OT, Taghert PH (2009) RNA-interference knockdown of Drosophila pigment dispersing factor in neuronal subsets: the anatomical basis of a neuropeptide’s circadian functions. PLoS One 4: e8298

Sharma VK, Chandrashekaran MK (1998) Age-dependent modulation of circadian parameters in the field mouse Mus booduga. J Exp Zool 280: 321–326

Small GW, Ercoli LM, Silverman DH, Huang SC, Komo S, Bookheimer SY, Lavretsky H, Miller K, Siddarth P, Rasgon NL et al (2000) Cerebral metabolic and cognitive decline in persons at genetic risk for Alzheimer’s disease. Proc Natl Acad Sci U S A 97: 6037–6042

Stevanovic K, Yunus A, Joly-Amado A, Gordon M, Morgan D, Gulick D, Gamsby J (2017) Disruption of normal circadian clock function in a mouse model of tauopathy. Exp Neurol 294: 58–67

Stopa EG, Volicer L, Kuo-Leblanc V, Harper D, Lathi D, Tate B, Satlin A (1999) Pathologic evaluation of the human suprachiasmatic nucleus in severe dementia. J Neuropathol Exp Neurol 58: 29–39

Tranah GJ, Blackwell T, Stone KL, Ancoli-Israel S, Paudel ML, Ensrud KE, Cauley JA, Redline S, Hillier TA, Cummings SR et al (2011) Circadian activity rhythms and risk of incident dementia and mild cognitive impairment in older women. Ann Neurol 70: 722–732

Umezaki Y, Yoshii T, Kawaguchi T, Helfrich-Forster C, Tomioka K (2012) Pigment-dispersing factor is involved in age-dependent rhythm changes in Drosophila melanogaster. J Biol Rhythms 27: 423–432

Valentinuzzi VS, Scarbrough K, Takahashi JS, Turek FW (1997) Effects of aging on the circadian rhythm of wheel-running activity in C57BL/6 mice. Am J Physiol 273: R1957–1964

van Gool WA, Witting W, Mirmiran M (1987) Age-related changes in circadian sleep-wakefulness rhythms in male rats isolated from time cues. Brain Res 413: 384–387

van Wamelen DJ, Aziz NA, Anink JJ, van Steenhoven R, Angeloni D, Fraschini F, Jockers R, Roos RA, Swaab DF (2013) Suprachiasmatic nucleus neuropeptide expression in patients with Huntington’s Disease. Sleep 36: 117–125

Wang X, Wang L, Yu Q, Xu Y, Zhang L, Zhao X, Cao X, Li Y, Li L (2016) Alterations in the expression of Per1 and Per2 induced by Abeta31-35 in the suprachiasmatic nucleus, hippocampus, and heart of C57BL/6 mouse. Brain Res 1642: 51–58

Weitzman ED, Moline ML, Czeisler CA, Zimmerman JC (1982) Chronobiology of aging: temperature, sleep-wake rhythms and entrainment. Neurobiol Aging 3: 299–309

Xu Y, Wei X, Liu X, Liao J, Lin J, Zhu C, Meng X, Xie D, Chao D, Fenoy AJ et al (2015) Low Cerebral Glucose Metabolism: A Potential Predictor for the Severity of Vascular Parkinsonism and Parkinson’s Disease. Aging Dis 6: 426–436

Xue Y, Chiu JC, Zhang Y (2019) SUR-8 interacts with PP1-87B to stabilize PERIOD and regulate circadian rhythms in Drosophila. PLoS Genet 15: e1008475

Yang W, Xia Y, Hawke D, Li X, Liang J, Xing D, Aldape K, Hunter T, Alfred Yung WK, Lu Z (2012) PKM2 phosphorylates histone H3 and promotes gene transcription and tumorigenesis. Cell 150: 685–696

Zee PC, Rosenberg RS, Turek FW (1992) Effects of aging on entrainment and rate of resynchronization of circadian locomotor activity. Am J Physiol 263: R1099–1103

Zhao X, Zhao L, Yang H, Li J, Min X, Yang F, Liu J, Huang G (2018) Pyruvate kinase M2 interacts with nuclear sterol regulatory element-binding protein 1a and thereby activates lipogenesis and cell proliferation in hepatocellular carcinoma. J Biol Chem 293: 6623–6634

Zhou JN, Hofman MA, Swaab DF (1995) VIP neurons in the human SCN in relation to sex, age, and Alzheimer’s disease. Neurobiol Aging 16: 571–576

Zilberter Y, Zilberter M (2017) The vicious circle of hypometabolism in neurodegenerative diseases: Ways and mechanisms of metabolic correction. J Neurosci Res 95: 2217–2235

